# FRET assay for live-cell high-throughput screening of the cardiac SERCA pump yields multiple classes of small-molecule allosteric modulators

**DOI:** 10.1101/2023.02.22.529557

**Authors:** Osha Roopnarine, Samantha L. Yuen, Andrew R. Thompson, Lauren N. Roelike, Robyn T. Rebbeck, Philip A. Bidwell, Courtney C. Aldrich, Razvan L. Cornea, David D. Thomas

## Abstract

We have used FRET-based biosensors in live cells, in a robust high-throughput screening (HTS) platform, to identify small-molecules that alter the structure and activity of the cardiac sarco/endoplasmic reticulum calcium ATPase (SERCA2a). Our primary aim is to discover drug-like small-molecule activators that improve SERCA’s function for the treatment of heart failure. We have previously demonstrated the use of an intramolecular FRET biosensor, based on human SERCA2a, by screening a small validation library using novel microplate readers that can detect the fluorescence lifetime or emission spectrum with high speed, precision, and resolution. Here we report results from a 50,000-compound screen using the same biosensor, with hit compounds functionally evaluated using Ca^2+^-ATPase and Ca^2+^-transport assays. We focused on 18 hit compounds, from which we identified eight structurally unique compounds and four compound classes as SERCA modulators, approximately half of which are activators and half are inhibitors. While both activators and inhibitors have therapeutic potential, the activators establish the basis for future testing in heart disease models and lead development, toward pharmaceutical therapy for heart failure.

## Introduction

Sarco/endoplasmic reticulum calcium ATPase (SERCA), integrated in the sarcoplasmic reticulum (SR, muscle cells) or endoplasmic reticulum (ER, non-muscle cells) membrane in most mammalian cells, is integral for using Ca^2+^-dependent hydrolysis of ATP to fuel active transport of cytosolic Ca^2+^ into the SR or ER. In muscle, the activity of SERCA1a (skeletal isoform) or SERCA2a (cardiac isoform) is essential for relaxation (diastole), restoring SR Ca^2+^ following its release via Ca^2+^ channels (ryanodine receptors, RyR) for muscle contraction (systole). Decreased SERCA activity and excessive RyR leak results in failure to maintain the high gradient of [Ca^2+^] between the cytoplasm (mM) and the SR (sub-μM) during diastole (muscle relaxation) and are associated with heart failure (HF) in human and animal models^1^. Decreased SERCA activity has been attributed to multiple factors, including reduced SERCA gene expression, increased post-translational modifications, and altered interaction with regulatory proteins^1^. Overall, decreased SERCA activity and increased Ca^2+^-leak lead to a pathophysiological state of the cardiac myocyte^2^ (HF, cardiac hypertrophy, diabetic hypertrophy), skeletal myocyte (Brody’s disease and myotonic dystrophy) and non-muscle cells (Darier’s disease, diabetes, Alzheimer’s disease)^3^. Altered SERCA interaction with regulatory proteins (regulins), such as phospholamban (PLB), have been linked to ventricular dysfunction and maladaptive remodeling in failing hearts^4^. Of the seven regulins discovered^5^, the dwarf open reading frame (DWORF) regulatory peptide is the only one known to activate muscle-specific SERCA activity, both by direct activation of SERCA^6,7^ and by competing with PLB binding^8,9,^ and to prevent HF in a mouse model of dilated cardiomyopathy^10^. Current therapeutic measures include beta-blockers, angiotensin-converting enzyme (ACE) inhibitors, angiotensin-receptor blockers (ARB), and mineralocorticoid receptor antagonists. However, these do not directly target proteins responsible for dysfunctional Ca^2+^ cycling. Discovery of small molecules (potential drugs) that target specific proteins is needed to exert improved control measures for positive therapeutic outcomes.

In the present study, we seek primarily SERCA2a activators to alleviate heart failure^11,12.^ However, SERCA uncouplers are also of potential interest in other indications such as in nonshivering thermogenesis, enhancing metabolism and thus reducing obesity^13,14.^ Targeted SERCA uncouplers or inhibitors may be useful for treatment of cancer or malaria^15^.

SERCA2a is a large transmembrane protein, with the phosphorylation (P) and nucleotide-binding (N) domains forming the catalytic site, coupled by the actuator (A) domain (Fig. 1A). Large (5-10 nm) relative movements of these domains are coupled to Ca^2+^ transport, as detected in living cells by an intramolecular FRET biosensor (Fig. 1A)^16^. The interaction of small molecules with SERCA can induce measurable structural changes, detectable by this biosensor, that often correlate with function, making this biosensor a powerful tool for HTS discovery of SERCA-binding compounds^16,17.^

**Figure 1.**
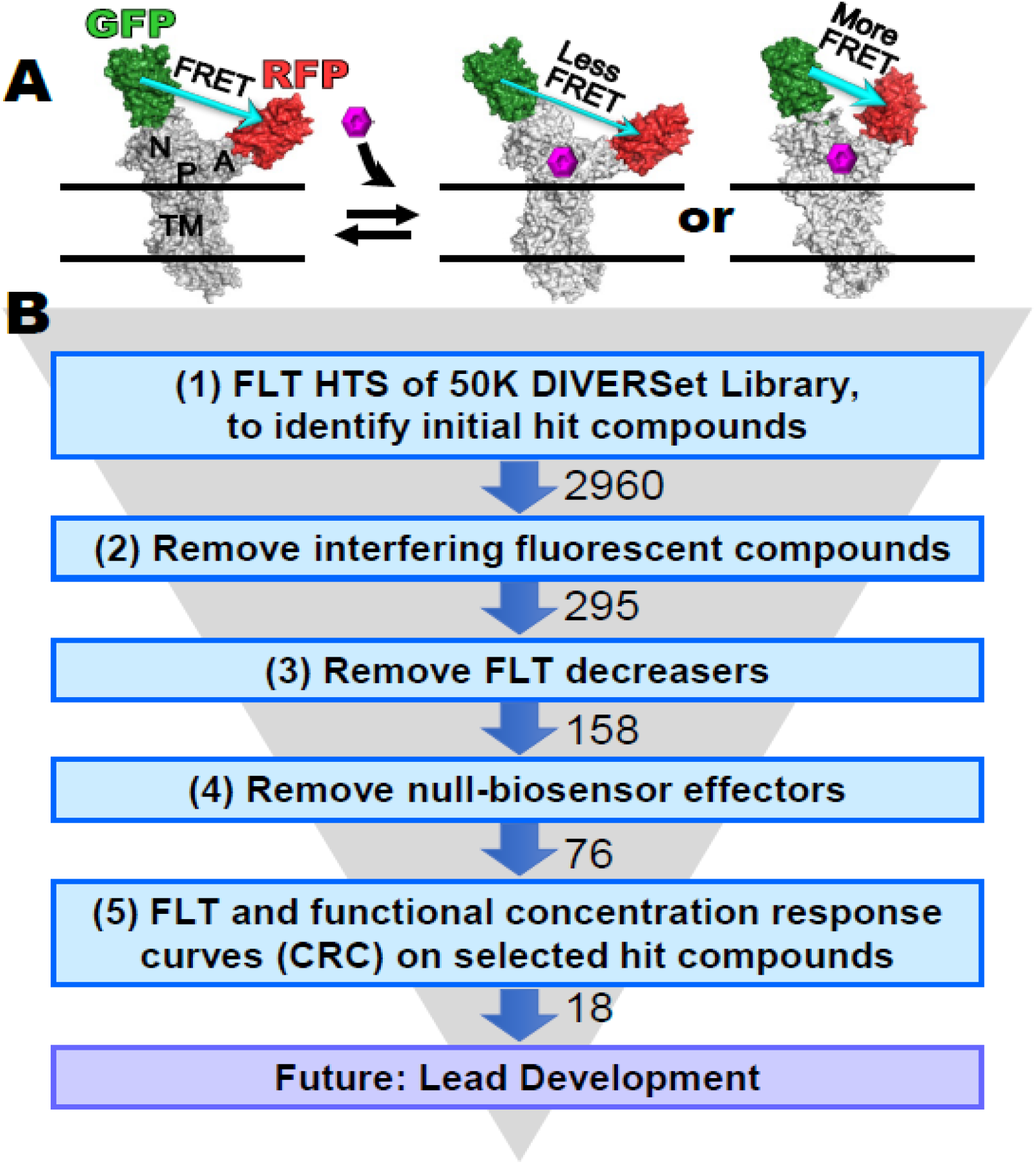
**(A)** FRET biosensor, human two-color SERCA2a (2CS), showing SERCA domains: nucleotide binding (N), phosphorylation (P), actuator (A) and transmembrane (TM). GFP is fused to N and RFP is fused to A. ΔFRET, measured from ΔFLT (fluorescence lifetime), is used to detect SERCA structural changes (**B**) Screening funnel describing the 5-step process undertaken in this study, involving measurements of FLT and SERCA function, with SERCA in live mammalian cells and in isolated pig cardiac SR membranes, respectively. (1) FLT changes caused by test compounds were measured using the target SERCA-specific FRET biosensor 2CS, to identify initial hit compounds. False hits were ruled out as compounds that (2) are fluorescent or affect the donor directly, (3) decrease FLT, or (4) affect FRET in a null biosensor, in which donor and acceptor are separated by a non-functional flexible peptide. (5) Concentration-dependence of FLT and SERCA function was measured to further prioritize hit compounds that could result in future lead development. Experimental details are provided in Methods.

In several previous early-stage drug discovery campaigns, we have focused on the SERCA regulator phospholamban (PLB)^18,19,^ but we have also tested and validated FRET biosensor constructs of SERCA alone^16,17,20,^ to detect binding of drug-like small-molecule compounds directly to SERCA. For this, we engineered a “two-color” SERCA (2CS, Fig. 1A) construct with fused eGFP and tagRFP fluorescent protein tags to the cytoplasmic N- and A-domains of SERCA, which detect relative motions of these domains during the enzymatic cycle responsible for Ca^2+^-transport^17,20,21.^ Using the intramolecular FRET measurement of 2CS, stably expressed in HEK293 cells, we previously validated this biosensor using the NIH Clinical Collection (NCC, 727 compounds)^17^ and the LOPAC library (1280 compounds)^20^. Along with the fluorescent biosensors, high-throughput is enabled by a FLT-PR (fluorescence lifetime plate reader) that scans a 1536-well plate with unprecedented precision and speed, determining FRET with 0.3% CV (30 times better than conventional intensity detection) in 2.5 min^18,22,23,^ enabling a high-precision screen of 50,000-compounds in two days. This instrument is equipped with simultaneous FLT detection at two emission wavelengths (two-channel lifetime detection)^21^, a function used to filter out fluorescent compounds that would be falsely identified as hit compounds. We have demonstrated the additional high throughput acquisition of fluorescence emission spectra using a spectral unmixing plate reader (SUPR), which helps detect (and rule out) interfering compounds, based on changes in the line shape in the donor-only region of the spectrum^21^. As in our previous projects involving other drug targets, the next logical step is to screen a 50,000-compound DIVERSet library, a diverse collection of drug-like small molecules that has yielded effective hit compounds in previous drug discovery projects^22-24^.

Here we apply our HTS platform using FRET lifetime measurements of two-color human cardiac SERCA (2CS) in living cells (Fig. 1B) to identify hit compounds. To validate selected hit compounds, we then acquire concentration response curves (CRCs) using both FRET and functional assays (Ca^2+^-ATPase activity and Ca^2+^-transport). We hypothesize that the combination of using improved fluorescence technology and screening a larger compound library (50K DIVERSet) will result in a larger and more diverse collection of hit compounds that more effectively regulate cardiac SERCA function, thus increasing the potential for discovering lead compounds for new heart failure therapeutics.

## Results

### FLT HTS of 50K DIVERSet library

2CS expressed in live HEK-293 cells was incubated with 5 nL of compound at a final [compound] of 10μM (from a DIVERSet library of 50,000 compounds) or DMSO (as a control), preloaded on 40 assay plates for 20 min prior to being read on the FLT-PR. FLT measurements were observed to be normally distributed, with a coefficient of variation (CV) of 0.4% across all 40 plates (Fig.2A). Plate-by-plate CV varied by < 1% (Fig. 2A).

**Figure 2.**
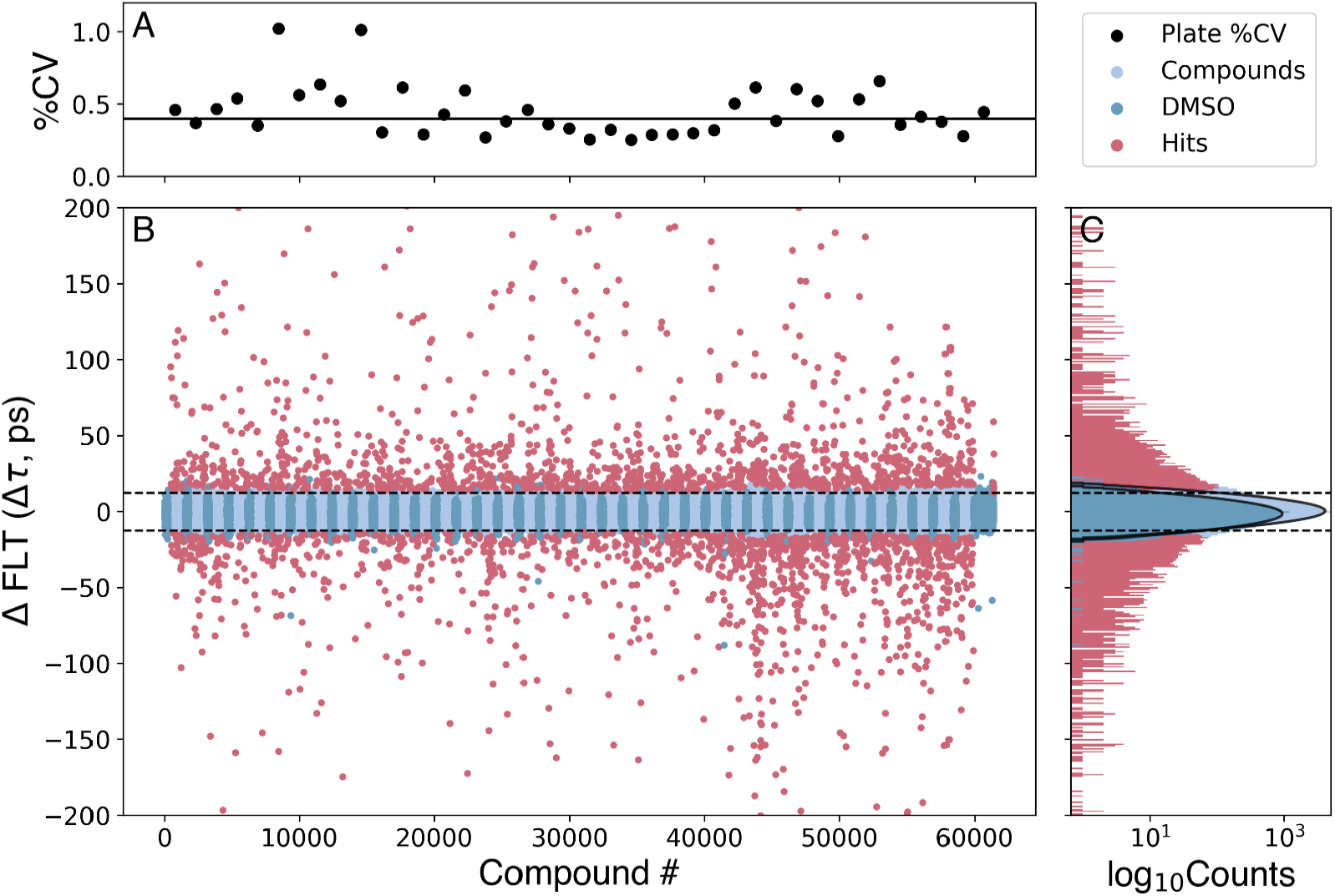
The human cardiac 2-color SERCA2a biosensor (2CS) in HEK293 cells was used to screen the 50,000-compound DIVERSet library. (**A**) Screen precision was determined by computing % CV for each plate using DMSO control wells, with a median value of 0.4% across 40 plates. (**B**) Change in lifetime (Δτ) was computed to find potential hits (red points) with a hit threshold set at a robust *z*-score of 3, resulting in 2960 FLT hit compounds for triage with the SUPR instrument and two-channel lifetime detection. DMSO controls (dark blue) were grouped in the plot to better illustrate plate boundaries. (**C**) The histogram of compounds not affecting 2CS (light blue, 1ps bin width) are normally distributed, similar to that of DMSO alone as shown by a fit of the populations to Gaussian distributions, which are shown in log scale. The horizontal lines in B, C illustrate the approximate cutoffs used, though actual cutoffs were determined on a plate-by-plate basis.

Compounds that significantly altered the structure of 2CS were determined by computing the change in lifetime (Δτ) for each compound compared to DMSO control sample (2CS plus DMSO), and the magnitude of this change was compared to the normal statistical fluctuation of the biosensor by computing the robust z-score (Methods). The distinct FLT changes induced by the potential hit compounds (Fig. 2B, red) are illustrated by the normal distributions of compounds not affecting SERCA (Fig. 2C, blue) and the control (Fig. 2C, dark blue). A hit threshold was set at a robust *z*-score of ± 3, resulting in 2960 FLT hit compounds (Fig. 1B, step 1). Interference from fluorescent compounds was removed (Fig. 1B, step 2) using both the similarity index (SI, detected by SUPR)^17^ and two-channel lifetime detection^21^. We also eliminated compounds that affected the lifetime detected from a donor-only (1CS) sample; 295 compounds remained.

More FLT decreasers than increasers were found to fail these tests, in alignment with previous studies^18,20,21,25^. FLT increasers are more advantageous for two additional reasons: (a) They offer greater reproducibility between repeats of a screen^18,20,21^. (b) Most previously identified SERCA modulators have been shown to be FLT increasers^16,17,20^. Therefore, we prioritized the 158 FLT increasers (termed “hit compounds” (Fig. 1B, step 3) for follow-up retesting and CRC evaluation.

### FLT retests of select hits compounds with 2CS and null biosensor

158 hit compounds were retested using 2CS (Step 4 in Fig. 1B; see Fig. 3A and C) and a null biosensor construct (Step 4 in Fig. 1B; see Fig. 3B and D), which consisted of GFP and RFP connected by a 32-residue unstructured flexible linker peptide (G32R)^20^. The null biosensor was used to rule out compounds that alter FLT by directly binding to the fluorescent proteins. A plot of the change in lifetime (Δτ) vs. the ΔR/G ratio (Fig. 3C and D) shows that the 2CS hits had little to no effects on the null biosensor.

**Figure 3.**
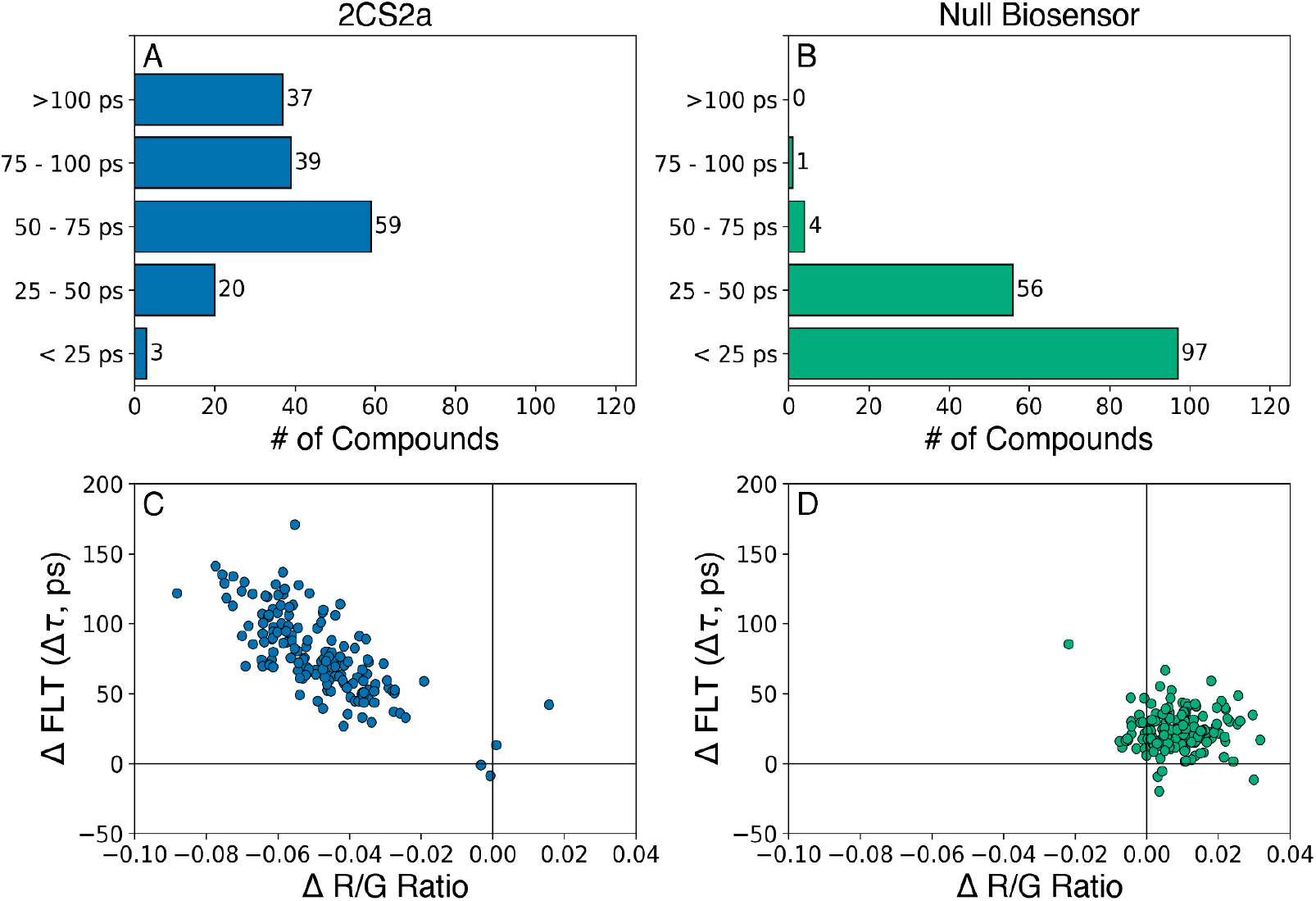
DIVERSet hit compound FRET retests. The most promising 158 hit compounds were selected (“cherry-picked”) and dispensed into 1536-well plates at 10 and 30 μM concentrations (n=3 wells for each drug concentration). Data is shown from the 30 μM screens for SERCA2a (A, C) and null biosensor (B, D). (**A**) Distribution of significant lifetime changes for the 2CS biosensor. (**B)** A null biosensor was counter-screened against the 158 hit compounds. Only five compounds displayed a lifetime change greater than 50 ps, indicating that our method for eliminating fluorescent compounds removes nearly all false positives. **(C**) and (**D**) Plots of the change in fluorescence lifetime vs the R/G ratio show excellent, reproducible correlation between the two techniques of lifetime (ΔFLT) and SUPR (ΔR/G) for the 2CS biosensor (**C**) and null biosensor (**D**).

Hit compounds that produced > 75 ps Δτ (76 compounds, Fig. 1B, step 4) (Fig. 3A) were targeted for further functional testing. We focused on 18 of these for the CRC testing, after the FLT data were subjected to the first four steps of the screening funnel (Fig. 1B) and compound repurchasing availability was determined. None of these compounds were in the PAINS (Pan-Assay INterference compoundS) category^26^, nor were they redox agents or metal chelators.

### Validation of hit compounds using FLT CRC

To further evaluate the 18 hit compounds, we determined the FLT response to compound concentrations ranging 0.78-100.μM (Step 5 in Fig. 1B). All 18 hit compounds (Table 1) decreased FRET (increased FLT) of the 2CS biosensor, suggesting that the compounds induced a structural change (Fig. 1, top) in the cytosolic headpiece region of SERCA2a. Compounds **1** and **4** showed a significant decrease in FRET at the lowest concentration, but no further effect at higher concentrations. The remaining 16 compounds decreased FRET with detectable EC_50_ (Fig. 4-7B, Table 1).

**Table 1:**
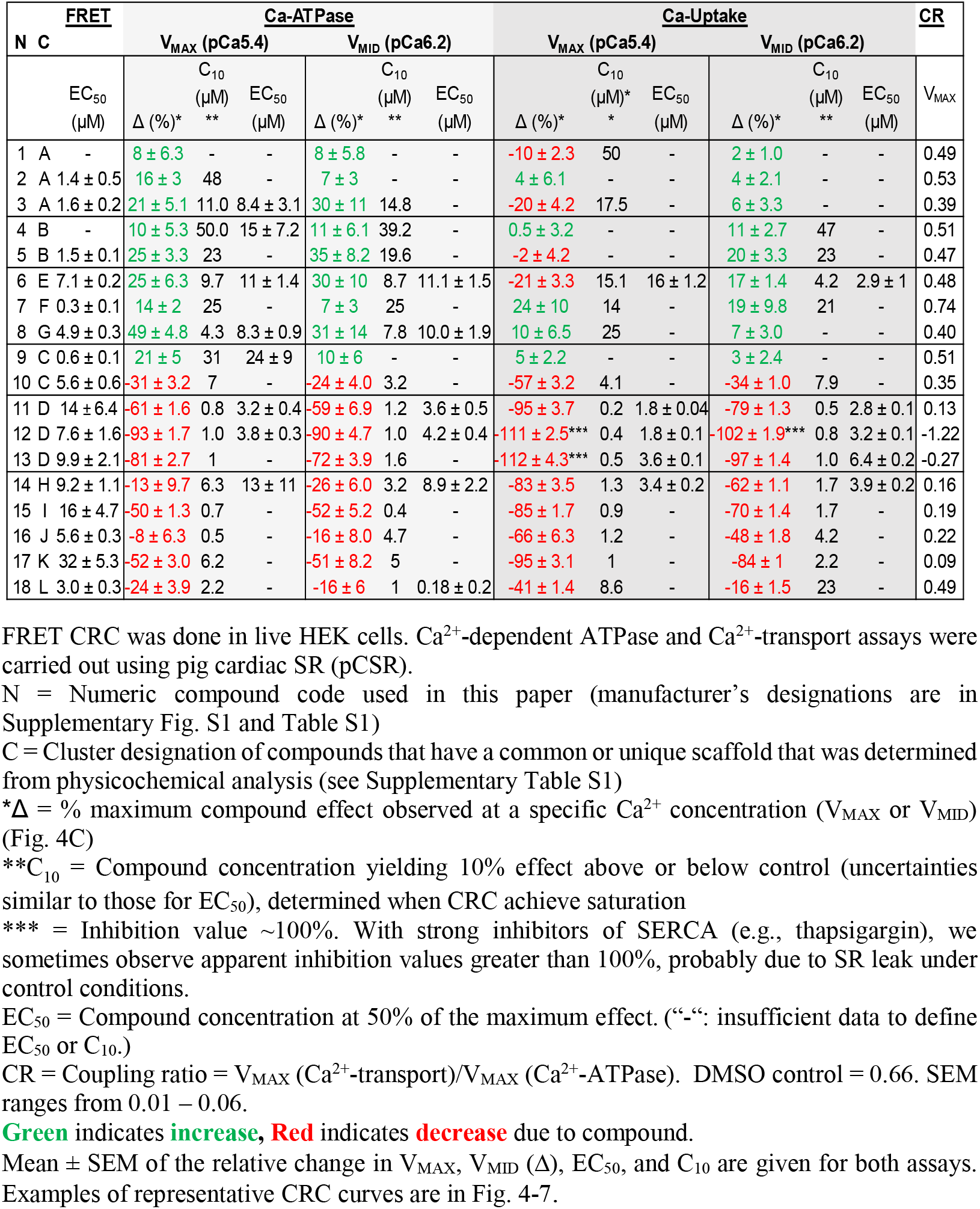
Results from the concentration response curves (CRC) for 18 hit compounds using FRET HTS, Ca^2+^-ATPase, and Ca^2+^ transport assays.

**Figure 4.**
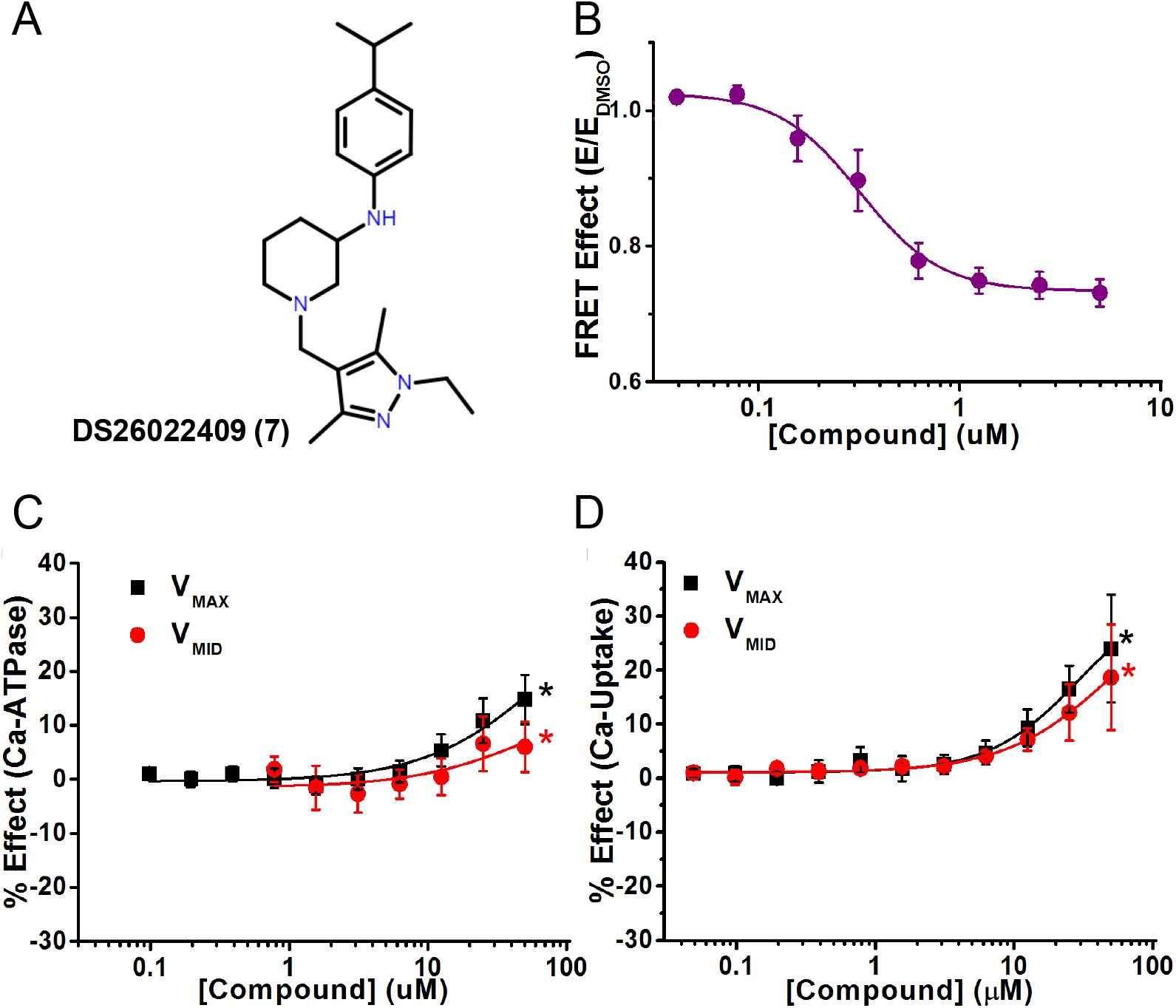
A representative activator compound of both Ca^2+^-ATPase and Ca-transport at saturating Ca^2+^ (V_MAX_). (**A**) Chemical structure of Compound DS26022409 (Compound **7**). (**B**) CRC of normalized FRET E in live HEK cells shows decreasing FRET response with increasing [compound]. (**C**) CRC of Ca^2+^-ATPase of SERCA2a in pCSR vesicles show activation at high Ca^2+^ (V_MAX_, black) and at midpoint Ca^2+^ (V_MID_, red). (**D**) CRC of Ca-transport shows activation at both V_MAX_ and V_MID_ Ca^2+^ (black and red, respectively). Data are presented as mean ± SEM, n = 3, *p<0.05.

### Effects of hit compounds on SERCA2a activity using Ca^2+^-ATPase and Ca^2+^-transport assays

To assess the impacts of FRET hit compounds on SERCA2a function, we used an absorbance-based enzyme-coupled NADH-linked Ca^2+^-dependent ATPase assay (referred to below as ATPase) and a fluorescence-based Ca^2+^-transport assay (referred to below as transport), using pig cardiac SR vesicles enriched for SERCA2a^18^ (Step 5 in Fig. 1B). Ca^2+^-ATPase and Ca^2+^-transport activities were measured at V_MAX_ (saturating, pCa 5.4) and V_MID_ (subsaturating, midpoint, pCa 6.2) [Ca^2+^], revealing activators (Compounds **1-9**) and inhibitors (Compounds **10-18**) (Table 1 and Supplementary Fig. S1), identified on the basis of potency (1/EC_50_) and efficacy (amplitude of the effect, Δ) (Table 1). Activators were compounds that increased Ca^2+^-ATPase and/or Ca^2+^-transport activities (at one or both [Ca^2+^]) (Table 1, Fig. 4-6). We grouped the activators in two subcategories based on the functional effects at the two Ca^2+^ concentrations: (1) activates both Ca^2+^-ATPase and Ca^2+^-transport (Compounds **2, 4, 7, 8**, and **9**), and (2) activates Ca^2+^-ATPase with divergent effects on Ca^2+^-transport (Compounds **1, 3, 5**, and **6**). We define “divergent” to indicate that the compound induces opposing effects at two different [Ca] (an increase at one [Ca] and a decrease at the other) in one assay. These effects indicate induced changes in the coupling ratio (CR), which is optimally two Ca^2+^ ions transported per molecule of ATP hydrolyzed^27-30^. This change was determined from the ratio of V_MAX_ (Ca^2+^-transport) to V_MAX_ (Ca^2+^-ATPase) (Table 1), as discussed below in **SERCA Activators**.

We identified inhibitors that induced strong (≥ 68%), moderate (34 to 67%), and mild (≤ 33%) inhibition of SERCA function. Therefore, we define four subcategories of compound inhibition: 1) strong inhibition of Ca^2+^-ATPase and Ca^2+^-transport (Compounds **11, 12, 13**), 2) moderate inhibition of Ca^2+^-ATPase and strong inhibition of Ca^2+^-transport (Compounds **15** and **17**), 3) mild inhibition of Ca^2+^-ATPase and moderate-to-strong inhibition of Ca^2+^-transport (Compounds **14** and **16)**, and 4) mild inhibition on Ca^2+^-ATPase and mild-to-moderate inhibition on Ca^2+^-transport (Compounds **10** and **18**) (Table 1, Fig. 7, and discussed below under **SERCA Inhibitors**).

### Classification of compounds by physicochemical characteristics

The 18 hit compounds were subjected to cheminformatic analysis, to determine whether any of the compounds shared a common chemical scaffold. Compounds with a Tanimoto coefficient and maximum common substructure (MCS)^31^ scores above 0.4 were binned as clusters, while those with scores below 0.4 were classified as singletons. The analysis yielded diverse scaffolds^31,32^ of the hit compounds (Supplementary Fig. S1 and Table S1). Four clusters of compounds with multiple examples (A-D in Table 1) were found, and the remaining eight were unique compounds (singletons) (E-L in Table 1 and Supplementary Fig. S1 and Table S1). The three compounds in cluster A (Compounds **1, 2**, and **3** in Table 1) have a common 5-(aryloxymethyl)oxazole-3-carboxamide)^33^, while those in cluster B (Compounds **4** and **5**) share a N-heteroaryl-N-alkylpiperazine. Cluster C (Compounds **9** and **10**) is defined by an amide linkage and Cluster D (Compounds **11, 12**, and **13**) by a piperidine scaffold. Clusters E-L (Compounds **6, 7, 8, 14, 15, 16, 17**, and **18**) contain a single compound (singleton) with no common scaffold with any other hit compound in this study. All of the hit compounds have physicochemical properties^34^, conducive of favorable drug disposition in vivo, including a low molecular weight (< 500), low cLogP (calculated partition coefficient for lipophilicity) values (< 5), low non-H rotatable bonds that describe the molecular flexibility (< 10), low degree of possible hydrogen bond formation (total number of hydrogen bond acceptors and donors should be less than 8), and low total polar surface area (tPSA < 140 Å) (Supplementary Table S1).

Next we describe in more detail the nine activators (Fig. 4-6) that are grouped into two subcategories, and the nine inhibitors (Fig. 7) that are grouped into three classes and four subcategories.

### SERCA Activators

In the first category of activators, Compounds **2, 4, 7, 8**, and **9** activated both Ca^2+^-ATPase and Ca^2+^-transport. Compound **7 (**Fig. 4A) (singleton F) decreased FRET of 2CS in live cells so that EC_50_ = 0.3 μM, indicating stabilization of the open conformation of SERCA, and accelerated Ca^2+^-ATPase to induce ΔV_MAX_ = 14% and ΔV_MID_ = 7% (Fig. 4C and Table 1). Compound **7** induced the highest increase in Ca^2+^-transport of all the compounds at both V_MAX_ (24%) and V_MID_ (19%), (Fig. 4D), which was greater than that of Ca^2+^-ATPase activity (Fig. 4D). The CR increased to 0.74 compared to control (0.66, Table 1). Saturation of CRC was not reached at the highest [compound] measured, so the functional EC_50_ was not determined, therefore we determined C_10_, the compound concentration that increases function by 10%. At V_MAX_ and V_MID_, C_10_ was 25 μM for Ca^2+^-ATPase, 14 μM and 21 μM for Ca^2+^-transport (Table 1). This compound will be placed at high priority for future optimization by medicinal chemistry and testing in animal models.

Compound **8** (singleton G) (Fig. 5A) decreased FRET (EC_50_ = 4.9 μM, Table 1) and increased Ca^2+^-ATPase at both V_MAX_ (49%, the largest increase observed in the screen) and V_MID_ (31%) (Table 1 and Fig. 5C). For Ca^2+^-transport, effects (Δ values) were lower (10% for V_MAX_ and 7% for V_MID_, Fig. 5D), decreasing CR to 0.4 (Table 1). EC_50_ values for Ca^2+^-ATPase were not significantly different at V_MAX_ and V_MID_, and were ~ 2x greater than the values observed by FRET. C_10_ was 4.3 μM (V_MAX_) and 7.8 μM (V_MID_), indicating significant ATPase activation at low dosage. C_10_ for Ca-transport was 25 μM at V_MAX_, and was not determined at V_MID_.

**Figure 5.**
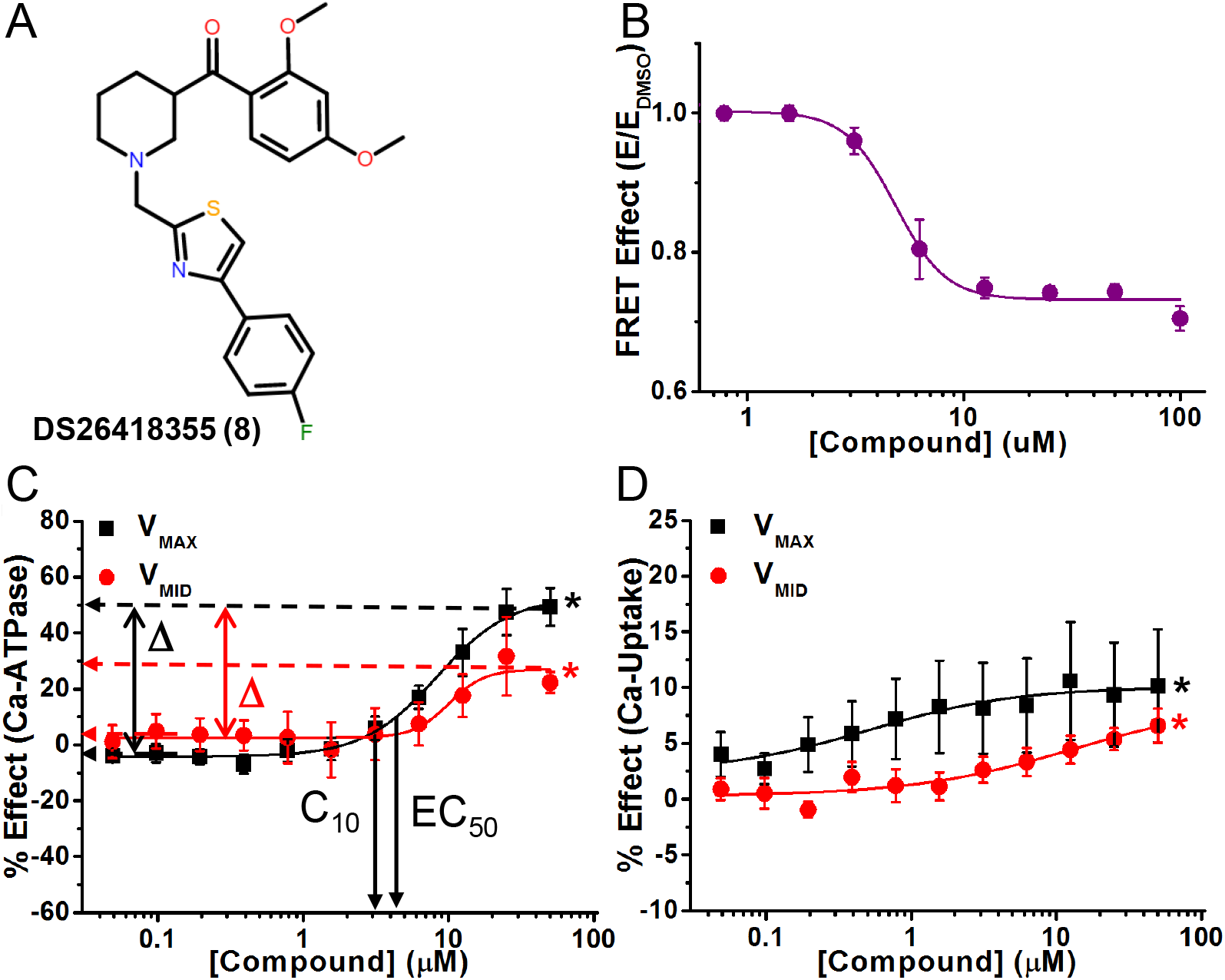
A representative activator compound that decreases FRET and uncouples Ca^2+^-ATPase from Ca^2+^-transport activity. (**A**) Chemical structure of Compound DS26418355 (Compound **8**). (**B**) CRC of normalized FRET E in 2CS biosensor in live HEK cells shows decreasing FRET response with increasing [compound]. (**C**) CRC shows Ca^2+^-ATPase activation in pCSR vesicles under both V_MAX_ (black) and V_MID_ (red) Ca^2+^. (**D**) Activation was less for Ca^2+^-transport of SERCA2a in pCSR vesicles. Mean ± SEM, n = 3, *p<0.05.

Compound **2 (**cluster A) induced similar Ca^2+^-ATPase activation (16 % at V_MAX,_ 7% at V_MID_) as observed with Compound **7**, but induced smaller increases on Ca^2+^-transport functions (Table 1), decreasing CR to 0.53. Compounds **4** (cluster B) and **9 (**cluster C) also showed similar effects as Compound **7**, increasing both activities. Compound **4** increased Ca^2+^-ATPase at both V_MAX_ and V_MID_ by ~10% and increased Ca^2+^-transport at both V_MAX_ (0.5%) and V_MID_ (11%). Compound **9** increased both V_MAX_ (21%) and V_MID_ (10%) for Ca^2+^-ATPase, with smaller increases in Ca^2+^-transport (3-5%). Compounds **4** and **9** decreased CR to similar extents (0.51) (Table 1).

In the second category of activators, Compounds **1** and **3** (cluster A), **5** (cluster B), and **6** (singleton E) activated Ca^2+^-ATPase at both V_MAX_ and V_MID_, but induced divergent effects on Ca^2+^-transport. Compound **6** (Fig. 6A) decreased FRET with EC_50_ = 7.1 μM (Fig. 6B, Table 1), while moderately activating Ca^2+^-ATPase activity by 25% at V_MAX_ and by 30% at V_MID_ (Fig. 6C), with EC_50_ = 11 μM for both V_MAX_ and V_MID_. It induced divergent effects during Ca^2+^-transport, inhibiting V_MAX_ by 21% and activating V_MID_ by 17% (Table 1 and Fig. 6D). C_10_ was similar at the two Ca^2+^ concentrations (9.7 μM and 8.7 μM), but was significantly different for Ca^2+^-transport (15 μM at V_MAX_, 4 μM at V_MID_). CR was decreased to 0.48 by Compound **6**.

**Figure 6.**
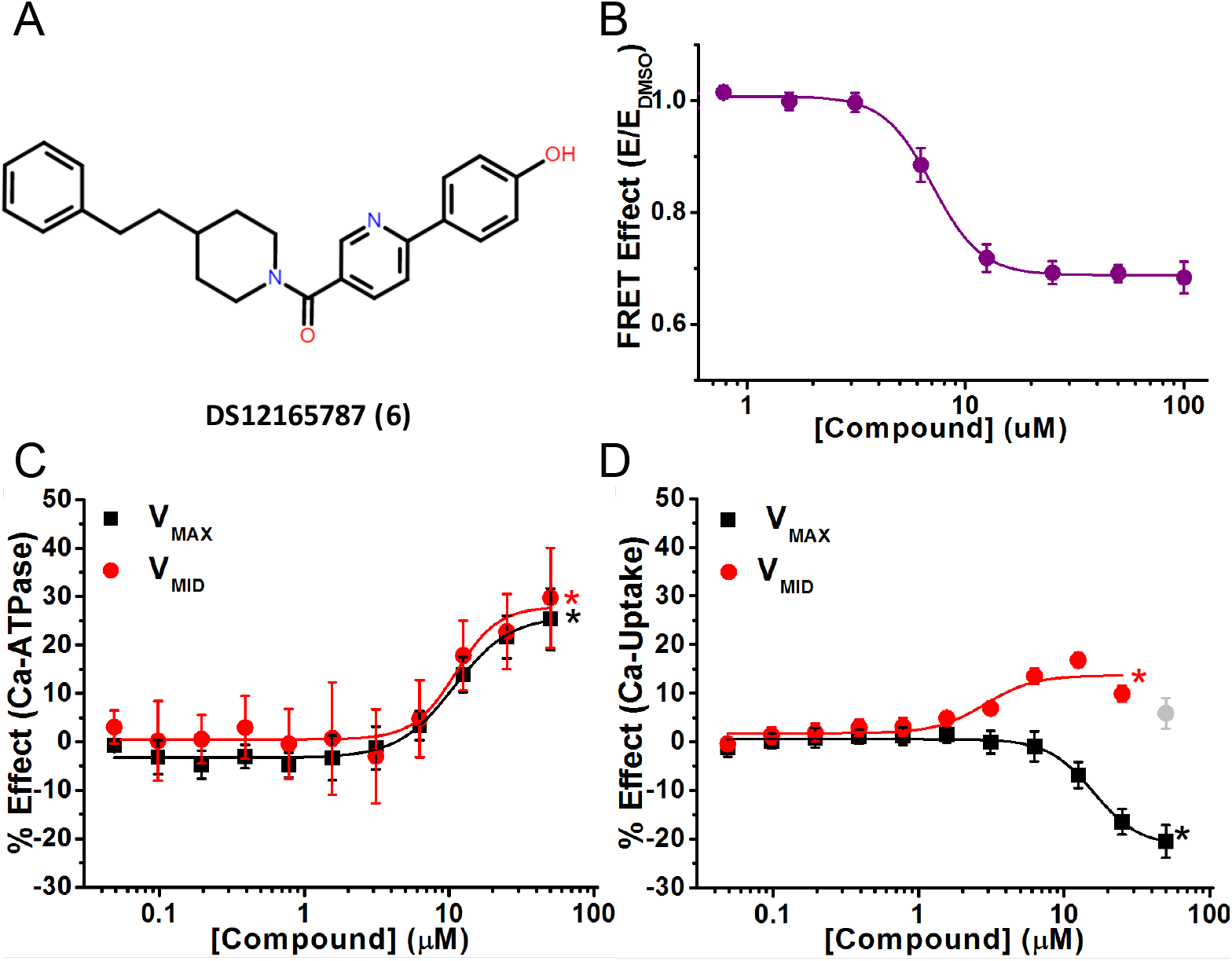
A representative activator compound that decreases FRET, increases Ca^2+^-ATPase activity, and has divergent effects on Ca^2+^-transport activity. (**A**) Chemical structure of Compound DS12165787 (Compound **6**). (**B**) Concentration response curve (CRC) of normalized FRET E shows decreased FRET with a half maximal effect (EC_50_) at 7.1 ± 0.2 μM. (**C**) CRC shows Ca^2+^-ATPase activation of SERCA2a in pig CSR vesicles at V_MAX_ (black, pCa 5.4) and V_MID_ (red, pCa 6.2) [Ca^2+^] (**D**) CRC of Ca^2+^-transport of SERCA2a in pig CSR vesicles, showing inhibition for V_MAX_ (black) and activation for V_MID_ (red). Grey data point was omitted from fitting. Δ, C_10_, and EC_50_ are reported in Table 1 for (C) and (D). Data is presented as mean ± SEM, n = 3, *p<0.05.

Compound **5** activated Ca^2+^-ATPase moderately at V_MAX_ (25%) and V_MID_ (35%). Ca^2+^-transport was inhibited slightly at V_MAX_ (2%), but activated at V_MID_ (20%) (Table 1). Compounds **1**, and **3** induced low activating effects at V_MID_ for Ca^2+^-transport (2% and 6%), but they inhibited Ca^2+^-transport at V_MAX_ (−10% and -20%) (Table 1). Compounds **1** and **5** induced similar decreases in the CR at V_MAX_ (0.0.49 and 0.47), while Compound **3** induced a slightly smaller CR of 0.39 (Table 1). These effects are similar to that of unphosphorylated phospholamban (PLB) in cardiac SR^35^.

### SERCA Inhibitors

Compounds **10**-**18** all decreased Ca^2+^-ATPase and Ca^2+^-transport activities at both V_MAX_ and V_MID_. Compared with FRET EC_50_ of the activators (0.3-7 μM), most of the inhibitors (Compounds **10**-**18**) showed weaker affinity, with FRET EC_50_ values in the range of 5-32 μM, but the maximum functional effects (efficacies) of the inhibitors tended to be greater (Table 1).

Compound **12** (cluster D) strongly inhibited both the Ca^2+^-ATPase and Ca^2+^-transport activities (Fig. 7C and D) to levels similar to the well-known SERCA inhibitor, thapsigargin, although thapsigargin acts with much greater affinity (EC_50_ ≈ 7.5 nM^18^) than compound **12** (EC_50_ = 3.8 μM) (Table 1).

**Figure 7.**
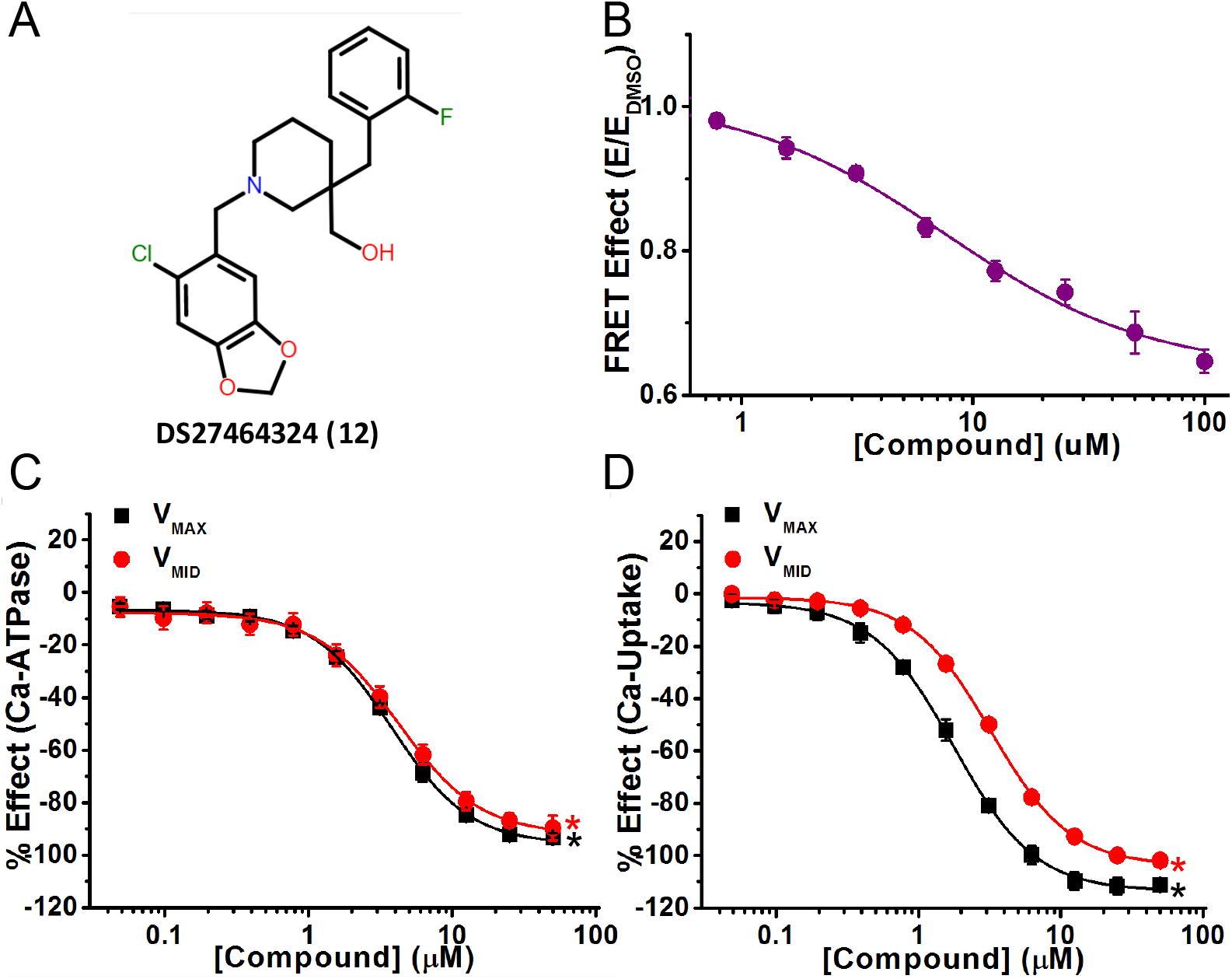
A representative inhibitor compound that severely decreases FRET and inhibits both Ca-ATPase and Ca-Transport activity. (**A**) Chemical structure of Compound DS27464324 (Compound **12**). (**B**) CRC of normalized FRET E shows decreased FRET response in 2CS biosensor. (**C**) CRC shows inhibition of Ca-ATPase in pCSR under both V_MAX_ (black) and V_MID_ (red) [Ca]. (**D**) CRC shows inhibition of Ca-transport in pCSR under V_MAX_ (black) and V_MID_ (red) [Ca]. Data is presented as mean ± SEM, n = 3, *p<0.05.

Compounds **11, 12**, and **13** (cluster D) showed similar inhibition of both SERCA2a functions: Ca^2+^-ATPase was inhibited by 61%, 93%, and 81% at V_MAX;_ by 59%, 90%, and 72% at V_MID_. Ca^2+^-transport was completely inhibited. Compared with Ca^2+^-ATPase, Ca^2+^-transport inhibition at V_MID_ required slightly higher compound concentration as shown by the shift to the right of the red curve (Fig.7D).

Compounds **15** (singleton I) and **17** (singleton K) induced moderate inhibition of both activities, decreasing V_MAX_ and V_MID_ by ~ 50% for Ca^2+^-ATPase and slightly more for Ca^2+^-transport (70-85%). C_10_ values for Compound **15** were ~1 µM or less, slightly higher for Compound **17** (5-6%) (Table 1) for Ca^2+^-ATPase, and similar for Ca^2+^-uptake at both V_MAX_ and V_MID_ ranging from (0.9-1.7%).

Compounds **14** (singleton H) and **16** (singleton J) induced mild inhibition of Ca^2+^-ATPase, but a considerably larger effect of moderate-to-strong inhibition of the Ca^2+^-transport. Ca^2+^-ATPase decreased by 13% and 8% at V_MAX_, 26% and 16% at V_MID_. Ca^2+^-transport was inhibited by 83% and 66% at V_MAX_, 62% and 48% at V_MID_. C_10_ values ranged from 0.5 to 7μM, for Ca^2+^-ATPase at V_MAX_ and V_MID_.

Compounds **10** (cluster C) and **18** (singleton L) induced mild inhibition of Ca2+-ATPase and moderate inhibition of Ca^2+^-transport. At V_MAX_, Ca^2+^-ATPase was inhibited by 31% and 24%, respectively; while at V_MID_ it was inhibited by 24 and 16%, respectively. At V_MAX_ Ca-transport was inhibited by 57% and 41%, and at V_MID_ by 34% and 16%. C_10_ values were 1 μM and 7 μM for Ca^2+^-ATPase, and 4.1 μM and 22.7 μM for Ca^2+^-transport.

## Discussion

We identified new compounds based on an increase in donor FLT, within a human cardiac 2CS biosensor expressed in live mammalian cells, indicating a decrease in FRET, implying that the actuator (A) and nucleotide-binding domains (N) of SERCAC2a moved further apart. We confirmed that these compounds affect SERCA activity using Ca^2+^-ATPase and Ca^2+^-transport assays with SERCA2a in native SR preparations, where we further categorized them as activators or inhibitors. We identified two subcategories of activators, whereby the compound either (1) activates both Ca^2+^-ATPase and Ca^2+^-transport activities (Compounds **2, 4, 7, 8**, and **9**) (Fig. 4 and 5) and (2) activates Ca^2+^-ATPase with divergent effects on Ca-transport (Compounds **1, 3, 5**, and **6**) (Fig. 6). We identified four subcategories of inhibitors based on the extent of Ca^2+^ATPase and Ca^2+^-transport decrease for cardiac SERCA2a (Table 1 and Fig. 7).

In general, the FRET EC_50_ values were smaller (indicating higher potencies) for activators (Compounds **1-9**; 0.3-7μM) than for inhibitors (Compounds **10-18**; (3-32μM) (Table 1). The potencies observed by FRET and function are not precisely correlated, probably because the assays were performed on different types of samples (live cells vs purified proteins), measuring different properties (structure vs function).

Functional CRC assays showed that inhibitors tended to induce larger changes (indicating higher efficacies) than activators, in both Ca^2+^-ATPase activity and Ca^2+^-uptake. Also, most inhibitors induce a larger change in Ca^2+^-uptake than in Ca^2+^-ATPase, decreasing the coupling ratio. In general, the C_10_ and EC_50_ values were smaller, indicating greater potency, for inhibitors than for activators.

Effects of most activators were to reduce the CR, as they induced larger changes in the Ca^2+^-ATPase than in corresponding the Ca^2+^-uptake assay. The most notable exception is Compound **7**, which activates Ca^2+^-transport even more than it activates Ca^2+^-ATPase, increasing CR. This compound will be a *high priority* as a lead compound for future efforts in medicinal chemistry and assays of physiological function.

Ten compounds were binned into four clusters (A-D), while eight compounds were classified as singleton (E-L) (Table 1). Many compounds showed similar functional traits, suggesting that there are ligand-sensing sites in SERCA2a that recognize a range of scaffolds, or that the ligand-binding sites are close to each other, providing potentially powerful tools in the design of future compounds^36,37,38^. Compounds **2** (cluster A), **4** (cluster B), **7** (singleton F), **8** (singleton G), and **9** (cluster C) induced similar effects of moderate activation on V_MAX_ in the Ca^2+^-ATPase assay with smaller activating effects on Ca^2+^-transport. Compounds in activator clusters A (Compounds **1** and **3**) and B (Compound **5**) along with Compound **6** from singleton E, showed similar functional effects: moderate activation of V_MAX_, with smaller activation of Ca^2+^-transport (decreased coupling) at both [Ca^2+^]. The compounds in clusters D (**11, 12**, and **13**), C (**10**), and H-L (**14**-**18**) of inhibitors also induced similar functional effects.

There was little or no overlap in the hit compounds identified in our previous FRET HTS screen of the same (DIVERSet) library with another biosensor for tumor necrosis factor receptor 1 (TNFR1^22^. There was 81% overlap in the fluorescent compounds detected in these two HTS screens, indicating that our FRET HTS screening methodologies independently and successfully removed the fluorescently interfering compounds^21,24.^ In another HTS study of the DIVERSet library, using a SERCA functional (Ca^2+^-ATPase) assay in the primary screen, we discovered several activators, several of which showed isoform specificity for either SERCA1a or SERCA2a^28^. However, there was negligible overlap between hit compounds identified in that ATPase HTS study and in the current study that used FRET in the primary screen. This observation highlights the value of complementary HTS assays for the same target.

SERCA activators are needed when cardiac relaxation is impaired, as in early-onset diastolic dysfunction that precedes systolic impairment in HF^1^, diabetic cardiomyopathy^39^, Alzheimer’s disease^40^, or Duchenne muscular dystrophy (DMD)^41^. The stimulation of SERCA2a activity, as a novel therapeutic measure to relieve cardiac dysfunction in heart failure without arrhythmogenic effects, is a promising strategy to be used in combination with other first-line therapeutic agents such as β-blockers and ACE inhibitors^42^. Until recently, only a few compounds were known to stimulate SERCA: CDN1163 (stimulates Ca^2+^ transport)^43,19,^ CP-154526 (increases the apparent Ca^2+^ affinity of SERC2a)^44^, Ro 41-0960 (increases SERCA maximal activity in high Ca^2+^)^44^, and istaroxime (stimulates SERCA activity)^45^. However, our recent HTS using Ca^2+^-ATPase activity as the target HTS assay identified ~19 new activators of SERCA^28^, and we identified nine activator compounds in the present study. A SERCA activator from our previous work shows promise as a therapeutic target for Alzheimer’s disease, as it rescued memory function in a mouse model for the disease^40^ as well as for DMD as it was shown to ameliorate dystrophic phenotypes in dystrophin-deficient mdx mice^41^. Of all these SERCA2a activators only istaroxime, a known Na^+^/K^+^ transporting ATPase inhibitor as well as an inotropic/lusitropic agent acting to enhance SERCA2a activity, has been in phase IIb clinical trials for treatment of heart failure^45,46.^ However, because of its unsuitability for human usage (poor gastrointestinal absorption, high clearance rate, and extensive metabolic transformation)^46^, istaroxime derivatives were designed from QSAR studies and a new promising class of SERCA2a activators has been identified^47,48,49^.

Compounds **1, 3, 5**, and **6** induced small effects on the V_MAX_ of Ca^2+^-ATPase (~10-25% increase) and induced a negative effect on the V_MAX_ of Ca^2+^-transport (Fig. 6C and D), thus decreasing the CR, which is likely to increase heat output^13,52,54^. These effects are similar to that of SLN on SERCA1a (skeletal muscle), where SLN induces no observable effect on the V_MAX_ of Ca-ATPase but reduces the V_MAX_ of Ca^2+^-transport (SERCA1a uncoupling), thus reducing CR^50^. This results in futile cycling of SERCA1a and higher usage of ATP, resulting in increased non-shivering thermogenesis (NST)^50^. Another contributor to NST is Ca^2+^ leak from SR to sarcoplasm through RyR channels, stimulating SERCA to re-sequester Ca^2+^ into the SR, thus using more ATP and generating heat^51^. This increase in energy expenditure in muscle has been suggested as a potential therapeutic strategy for weight loss^50,52.^ Thus, the SERCA uncouplers in this study may serve as the basis for further drug development targeting weight loss.

Over the past several decades (~ 60 years), research on the potential for small-molecule SERCA inhibitors as oncology therapeutics has yielded hundreds of SERCA inhibitors with varying potencies and efficacies^15^. Similarly, our discovery of new SERCA inhibitors with a range of potencies and efficacies is likely to be advantageous for treatment of non-cardiac applications^15,53.^

In the present study, the 2CS biosensor has been used to identify novel small-molecule effectors of SERCA that have diverse chemical scaffolds for binding to SERCA, resulting in an array of hit compounds that are activators and inhibitors. Most importantly, we discovered a potential lead compound (Compound **7**) that activates Ca^2+^-uptake more than the Ca^2+^-ATPase, increasing the CR, so this will be a *high priority* for future efforts in medicinal chemistry and assays of physiological function. The enabling technology included three novel plate-readers, the FLT instrument used in the primary screen, and two spectral instruments that were used to remove interference of fluorescent compounds, allowing us to focus on valid SERCA activators and inhibitors. In the future, hits from the present study will be evaluated in more functional detail, including studies on multiple SERCA isoforms and on intact muscles and animals. Medicinal chemistry will be applied to elucidate structure-activity relationships, with the goal of designing analogs with greater potency and specificity^22,54.^ We will also expand our approach to much larger compound libraries, since our primary screening technology is capable of evaluating thousands of compounds per hour.

## Methods

### Molecular biology

A two-color intramolecular human SERCA2a (2CS) biosensor, based on human cardiac SERCA2a fused to green fluorescent protein (eGFP) and red fluorescent protein (tagRFP) was developed to detect structural changes that are related to the functional changes of SERCA^20^. Briefly, tagRFP was genetically fused to the N-terminus of SERCA and eGFP was inserted as an intrasequence tag before residue 509 in the nucleotide-binding domain (N-domain)^55,56.^ A donor-only (1CS) biosensor was created in a similar manner as the 2CS biosensor with the exception of the construct containing only eGFP. The fluorescent proteins fused to SERCA in 2CS and 1CS do not significantly affect SERCA activity, in membranes purified from HEK cells^18,20.^ A null biosensor construct consisting of eGFP and tagRFP connected by a 32-residue unstructured flexible linker peptide (G32R) was created as described previously^18,20.^ All constructs were cloned into expression vectors containing the genes for antibiotic resistance to G418, puromycin, or blasticidin.

### Cell culture

Stable cell lines were generated using either HEK293 (ATCC, Manassas, VA) or HEK293-6E (National Research Council, Canada) cells^20^. Briefly, cells were transiently transfected with 2CS, 1CS, or G32R null biosensor plasmids using Lipofecatime 3000 or 293fectin (Thermo Fisher Scientific). Flow cytometry was used to select and enrich for the population of cells expressing respective biosensors. Stable HEK293 cell lines were maintained in phenol red-free DMEM media (Gibco, Waltham, MA) supplemented with 2 mM GlutaMAX (Gibco, Waltham, MA), 10% fetal bovine serum (FBS) (Atlanta Biologicals, Lawrenceville, GA), 1 IU/mL penicillin/streptomycin (Gibco, Waltham, MA), and 250μg/mL G418 (Fisher Scientific). Stable HEK293-6E cell lines were maintained in F17 media (Sigma Aldrich) supplemented with Kolliphor p188 (Sigma Aldrich, St. Louis, MO), 200 nM/mL GlutaMAX, and either 1μg/mL puromycin (Invitrogen, Carlsbad, CA) or 2μg/mL blasticidin (Goldbio) as a selection antibiotic. All cell lines were grown at 37°C with 5% CO_2_.

### Compound handling

A DIVERSet 50,000 compound library was purchased from ChemBridge Corporation (San Diego, CA) at a 10 mM stock concentration for each compound. All compounds met the high quality standard of 100% identification by NMR and/or LC-MS and have a minimum purity of 85% and their identity verified using LC-MS/ELSD as confirmed by the ChemBridge Corporation. For the FRET HTS initial screens, the compound library was reformatted into 384 well Echo compatible plates using the Biomek FX (Beckman Coulter, Miami, FL) and 5 nL of either compound (columns 3-22 and 27-46) or DMSO (columns 1-2, 23-26, and 47-48) was dispensed into 1536 well black polystyrene assay plates (Greiner, Kremsmünste, Austria) using an Echo 550 liquid dispenser (Beckman Coulter) to yield a final assay screening concentration of 10μM. The low autofluorescence and low interwell cross-talk of these plates made them advantageous for FRET measurements. Plates were heat sealed with a PlateLoc Thermal Microplate Sealer (Agilent, Santa Clara, CA) and stored at -20°C prior to use. The same methods were applied for subsequent FRET retesting of the reproducible hit compounds identified in the pilot screen, except that the [compound] was tested at 10μM and 30 μM in triplicate.

FRET CRC assay plates (0.78-100 μM compound range) with at least ten different compound concentrations were made by adding the appropriate volume of compound or DMSO into black 384 well plates (Greiner Bio-One) using a Mosquito HV (SPTLabTech, United Kingdom). Subsequent ATPase and Ca^2+^-transport CRC assay plates (0-50 μM compound range) with repurchased compounds were made in a similar manner using with the Echo 550 (Beckman Coulter) using either 384 well transparent plates (Greiner Bio-One) or black-walled plates with transparent bottoms (Greiner Bio-One), respectively.

### HTS sample preparation and FRET measurements

On each day of screening, cells were harvested, washed three times with PBS, and centrifuged at 300*g* for 5min. Cells were filtered using a 70µm cell strainer and diluted to 1-2 × 10^6^ cells/mL. Cell concentration and viability were assessed using the Cell countess (Invitrogen) and trypan blue assay. During assays, cells were constantly and gently stirred using a magnetic stir bar at room temperature, keeping the cells in suspension and evenly distributed to avoid clumping. Cells were dispensed at 5μL or 50μL per well into assay plates (dispensed into 40 assay plates, each containing 1536 wells) pre-plated with either compound or DMSO using a Multdrop Combi liquid dispenser (Thermo Fisher Scientific, Pittsburg, PA) and sealed until needed. Because the kinetics of membrane permeability, diffusion, and/or binding of the compound to live cells may be compound-dependent, we tested two incubation times, 20 min and 120 min, for the FLT CRC. FRET EC_50_ values determined from both incubations were similar, but the 120 min incubation yielded a more reproducible and sigmoidal curve. Plates containing eight-point concentration curves of three tool compounds (known SERCA effectors) were also included on the plates as positive controls for the HTS FRET assay. The FRET HTS screen was performed over two days with a custom HTS fluorescence lifetime plate reader (FLT-PR) and spectral plate reader (SUPR) provided by Photonic Pharma LLC (Minneapolis, MN)^18^.

The same methods were applied for subsequent FRET retesting of the reproducible hit compounds identified in the pilot screen, except that the compound tested at 10μM and 30 μM [compound]. 158 hit compounds were picked from the library master plates and reloaded onto new assay plates for retesting with 2CS and a null biosensor. Then 18 hit compounds were selected and purchased from ChemBridge Corporation to determine CRC from FRET, ATPase, and Ca-transport assays using at least ten different concentrations by repeatedly scanning the 1536-well plates.

### FRET HTS instrumentation and data analysis

A detailed description of the high-throughput fluorescence lifetime plate reader (FLT-PR) and spectral unmixing plate reader (SUPR), manufactured by Fluorescence Innovations Inc and provided by Photonic Pharma, LLC was described previously^18,21.^ Briefly, for lifetime measurement with the FLT-PR, the observed donor-fluorescence waveform, *F* (t) was fit by a convolution of the measured instrument response function (IRF) and a single-exponential decay to obtain the lifetime (τ) of the donor fluorophore^57^ in the absence (τ_D_) and presence (τ_DA_) of the acceptor as described in Equation (1):

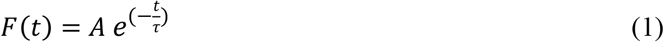

FRET efficiency (*E*) was determined as the fractional decrease of donor FLT in the absence and in the presence of acceptor as in Equation (2):

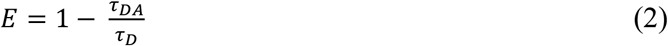

*E* was determined in the presence and absence of compound and normalized relative to *E* of the DMSO control. For spectral detection with the SUPR, the observed fluorescence emission spectrum F(λ) was fit by least-squares minimization of a linear combination of component spectra for donor (D), acceptor (A), cellular autofluorescence (C) and water Raman (W) as described previously^17^.

### HTS data analysis

FLT-PR data was used as the primary metric for flagging potential hit compounds. After fitting waveforms with a single exponential decay to quantify donor lifetime, the change in fluorescence lifetime (Δ FLT) was computed by performing a moving average subtraction in the order the plate was scanned with a window size of half a plate row (24 columns). The reasons for this are twofold: 1) plate gradients are often observed due to heating of the digitizer during acquisition and 2) performing ΔFLT computations with DMSO controls alone can sometimes result in artifacts as a half of the DMSO wells are on the edge of plates, which occasionally exhibit artifacts due to processes needed for the preparation of the drug library being tested. As most compounds are likely to be non-hits, and therefore DMSO like, computation of a moving average is an effective alternative to solving both gradient issues and edge-effect distortion of the primary metric for hit selection, Δτ. As hit compounds from FLT-PR were to be further triaged with a secondary technique (using the spectral plate reader), a generous cutoff was set at a robust *z*-score of 3 on a plate-by-plate basis. The robust z-score was used, where the median (*M*) and median absolute deviation (*MAD*) are used in place of the mean and standard deviation (Equation 3), to best capture the most hits, as the standard z-score is more subject to strong outliers (compounds that fall outside of the defined upper and lower limits)^18^.

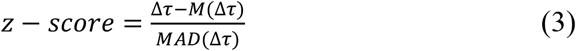

To remove “false positive” fluorescent compounds, the similarity index (SI^17^ was computed by comparing a region (500-540nm) of the donor only spectrum (*I*^(a)^) for each well to that of the plate-wide average DMSO spectrum (*I*^(b)^) in the same wavelength band as described in Equation 4^21^. Compounds that exceeded an SI robust z-score of 5 (corresponding to an SI of 2×10^−4^) were deemed likely fluorescent compounds and removed from consideration.

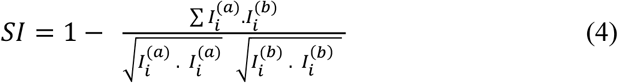

Spectral (SUPR) data was processed similarly to FLT-PR data, with the ΔR/G ratio being computed by applying the same moving average filter on the initial measurement of the ratio of the acceptor amplitude over the donor amplitude as found by fitting basis sets of the component spectra through least squares minimization. The hit threshold was also set using a robust z-score of 3. While the FLT-PR data and SUPR data showed a robust correlation, the FLT-PR data exhibited some strong outliers, presumably due to compounds directly modifying the donor lifetime. To eliminate these likely interfering compounds, correlation was enforced by eliminating compounds that exceed a robust z-score of 3 from the median value of the ratio of ΔFLT over the ΔR/G ratio metric. Additional interfering compounds were removed using two-channel lifetime detection^18^.

### Cardiac SR preparation

Cardiac SR vesicles were isolated from fresh porcine left ventricular tissue using differential centrifugation of the homogenized tissue as previously described ^58^. The SR vesicles were flash-frozen and stored at -80ºC until needed.

### Enzymatic SERCA activity assays of FRET hit compounds

Functional assays were performed using porcine cardiac SR vesicles^16^. An enzyme-coupled, NADH-linked ATPase assay was used to measure SERCA ATPase activity in 384-well microplates. Each well contained 50 mM MOPS (pH 7.0), 100 mM KCl, 1 mM EGTA, 0.2 mM NADH, 1 mM phosphoenol pyruvate, 10 IU/mL of pyruvate kinase, 10 IU/mL of lactate dehydrogenase, 7 µM of the calcium ionophore A23187 (Sigma), and CaCl_2_ was added to set free [Ca^2+^] to three different concentrations^59^. The Ca^2+^-ATPase were measured at V_MAX_ (saturating, pCa 5.4), V_MID_ (subsaturating, midpoint, pCa 6.2), and basal (non-activating, pCa 8.0) [Ca^2+^]. 10 μg/mL of SR vesicle, calcium, compound (0.048 to 50μM), and assay mix were incubated for 20 min at room temperature before measurement of functional assays with each of the 18 hit compounds, because a shorter incubation time than the FRET live-cell assays achieved optimal responses. The assay was started upon the addition of MgATP, at a final concentration of 5 mM (total volume to 80 μL), and absorbance was read in a SpectraMax Plus microplate spectrophotometer from Molecular Devices (Sunnyvale, CA) at 340nm.

### Ca^2+^-transport assays of FRET hit compounds

Ca^2+^-transport assays were performed with similar porcine SR samples as in the Ca^2+^-ATPase assays described above. The compound effect on the Ca^2+^-transport activity of SERCA2a was determined using an oxalate-supported assay in which the change in fluorescence in a Ca-sensitive dye, Fluor-4, was determined as previously described^18^. A buffered solution containing 50 mM MOPS (pH 7.0), 100 mM KCl, 30 mg/mL sucrose, 1 mM EGTA, 10 mM potassium oxalate, 2 μM Fluo-4, 30 µg/mL porcine cardiac SR vesicles, CaCl_2_ calculated to reach the free [Ca^2+^] (pCa 8.0, 6.2, and 5.4), and compound (0.048 to 50μM) was dispensed into 384-well black walled, transparent bottomed plates (Greiner Bio-One) containing the tested small molecule and incubated at 22ºC for 20 minutes while covered and protected from light. To start the reaction, MgATP was added to a final concentration of 5 mM, and the decrease in 485-nm excited fluorescence of Fluo-4 was monitored at 520 nm for 15 min using a FLIPR Tetra (Molecular Devices, San Jose, CA).

### Data analysis of FRET CRC assays of hit compounds

FRET efficiency (*E*) (Equation 2) was determined as the fractional decrease of donor (1CS) lifetimes (τ_D_) in the presence of acceptor (2CS) fluorophore (τ_DA_) due to FRET as described in Equation 1 and normalized to DMSO controls.

### Data analysis of Ca^2+^-ATPase and Ca^2+^-transport CRC assays

SERCA2a ATPase (or Ca-transport) activity at pCa 8.0 was subtracted from pCa 5.4 and pCa 6.2 values. The % effect ATPase (or Ca-transport transport) activity was normalized to the DMSO only (or 2CS in the absence of compound), and then were plotted against [Ca], and the curves were fitted using the Hill function, where *V* is the initial ATPase rate (or fluorescence rate), V_MAX_ is the ATPase (or Ca^2+^-transport) at saturating [Ca^2+^], and EC_50_ or *pK*_Ca_, or V_MID_ is the apparent Ca^2+^ dissociation constant as described previously ATPase (or Ca^2+^-transport) at (Midpoint Ca^2+^)^44^. These parameters and the [Ca^2+^] at 10% (*C*_10_) above or below baseline pCa (8.0) are reported in Table 1.

### Cheminformatic analysis of hit compounds

An online interactive program was used to perform cheminformatics analysis^60^ to determine whether the hit compounds had structural similarity by identifying common chemical scaffolds (core structural feature) using binning, multidimensional scaling (MDS), and compound similarity methods where the Tanimoto coefficient^31^ and maximum common substructure^31^ values were used to determine clustering (Supplementary Table S1). The physicochemical properties (for e.g. Lipinski Rule of 5) and bioactivity properties of the compounds were also used in the clustering analysis^34^. A cluster contained two or more compounds with similarity score > 0.4, while a unique compound with a similarity score < 0.4 was referred to as a singleton.

### Statistical analysis

Analysis of two-group comparisons was done by a two-tailed unpaired Student’s t-test (*p < 0.05) using the data analysis program Microsoft Excel (Santa Rosa, CA). Data are presented as mean ± SEM calculated from a minimum of three separate experiments (n = 3).

### Data availability

All the data discussed are presented within the article and Supplementary Information and are available from the corresponding authors (OR and DDT) on reasonable request.

## Acknowledgments

Prachi Bawaskar, Ben Grant, Simon J. Gruber, Evan W. Kleinboehl, Ang Li, Ji Li, Kurt Peterson, Seth L. Robia, and Tory M. Schaaf contributed to the conception of this project. Jesse E. McCaffrey, Bengt Svensson, Sarah Blakely Anderson, and J. Michael Autry provided helpful discussions. Marzena Brinkmann provided helpful advice on the chemical scaffold nomenclature. Fluorescence microscopy was performed at the UMN Imaging Center, flow cytometry at the UMN Lillehei Heart Institute, compound dispensing at the UMN Institute of Therapeutic Drug Discovery and Development, and spectroscopy at the UMN Biophysical Technology Center. This work was supported by NIH grants R01HL139065 (to DDT and RLC) and R37AG26160 (to DDT).

## Author contributions

DDT and RLC designed the research. SLY, ART, LNR, and PAB prepared samples and performed experiments. SLY, ART, and OR analyzed the data. OR wrote the first draft of the manuscript. OR, SLY, ART, RTR, CCA, RLC, and DDT edited and revised the manuscript.

## Additional information

### Supplementary Information

Available online

