## Supplemental Information for "FRET assay for live-cell high-throughput screening of the cardiac SERCA pump yields multiple classes of small-molecule allosteric modulators"

This document contains Supplementary Information for the Results section.

Page S2: Supplementary Figure S1 showing results from FRET HTS of DIVERSet library in Figure 2 and 3 showing 18 hit compounds.

Page S3: Supplementary Table 1 showing chemical names and physio-chemical properties of 18 hit compounds from Figure S1, resulting from analysis in Figures 2 and 3 in Results.

Page S5: Supplementary References

**Chemical Structures of selected hit compounds derived from the HTS FRET screening.**
The DIVERset (ChemBridge, San Diego, CA) 50,000 compound library was used to screen for HTS FRET structural changes in human two-color SERCA2a (2CS). Application of several analysis steps, outlined in the screening tree (shown in Fig. 1), on the fluorescence lifetimes in the presence of the compounds compared to that in the absence of compounds (Fig. 2 and 3) resulted in 18 hit compounds (Supplementary Fig. S1).

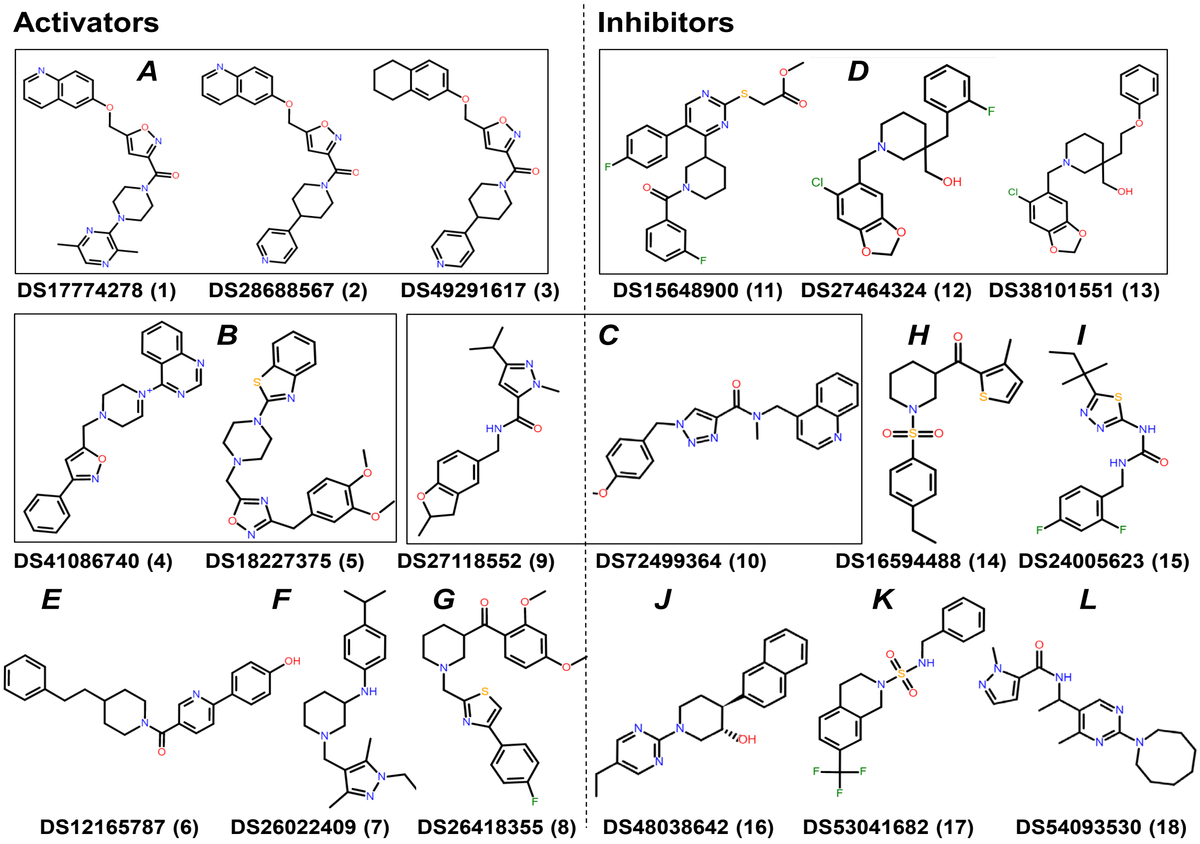


**Supplementary Figure S1.** Chemical structures of 18 hit compounds that decrease FRET *E* of 2CS by 3SD in HTS assay. The compound DIVERSet identification number (ID) and assigned numeric code (**1**-**18**) are listed below each compound. Compounds with a tanimoto coefficient [1] and maximum common substructure (MCS) score above 0.4 were binned in clusters in boxes (**A**-**D**) and as singleton structures (**E**-**L**).

**Supplementary Table S1: Chemical names and physicochemical properties of hit compounds**. The compound ID from the DIVERSet 50K library (ChemBridge, San Diego, CA) and assigned numeric code (**1-18**) are in the first column. Compounds with a tanimoto coefficient and maximum common substructure (MCS)[1] scores above 0.4 were binned as clusters (**A-D**), while those with scores below 0.4 were classed as singletons (**E-L**). The MSC or scaffold of the compound is in bolded text in its chemical name in 2nd column. The Lipinski's Rule of Five[2] physiochemical properties (MW, cLogP, rotatable bonds, hydrogen donors and acceptors, and total polar surface area) are listed for each compound.

| *Compound ID | Chemical Name | MW | cLogP | Rot. bonds | Hdon | HAcc | tPSA (Å) |
|---|---|---|---|---|---|---|---|
| | | **Activators** | | | | | |
| | | **A** | | | | | |
| DS17774278 (**1**) | 6-[(3-{[4-(3,6-dimethyl-2-pyrazinyl)-1-**piperazinyl]carbonyl}-5-isoxazolyl**)methoxy]quinoline | 444.5 | 3.17 | 5 | 0 | 7 | 97.5 |
| DS28688567 (**2**) | 6-[(3-{[4-(4-pyridinyl)-1-**piperidinyl]carbonyl}-5-isoxazoly**l)methoxy]quinoline | 414.5 | 4.15 | 5 | 0 | 6 | 81.4 |
| DS49291617 (**3**) | 4-[1-({5-[(5,6,7,8-tetrahydro-2-naphthalenyloxy)methyl]-3-**isoxazoly**l}**carbonyl)-4-piperidinyl**]pyridine | 417.5 | 4.5 | 5 | 0 | 0 | 68.4 |
| | | **B** | | | | | |
| DS41086740 (**4**) | 4-{4-[(3-phenyl-**5-isoxazolyl)methyl]-1-piperazinyl**}quinazoline | 371.4 | 3.6 | 4 | 0 | 5 | 58.3 |
| DS18227375 (**5**) | 2-(4-{[3-(3,4-dimethoxybenzyl)-**1,2,4-oxadiazol-5-yl]methyl**}**-1-piperazinyl**)-1,3-benzothiazole | 451.5 | 3.6 | 5 | 0 | 7 | 105 |
| | | **E** | | | | | |
| DS12165787 (**6**) | 4-(5-{[4-(2-phenylethyl)**piperidin-1-yl]carbonyl}pyridin-2-yl**)phenol | 386.5 | 4.88 | 5 | 1 | 3 | 53.4 |
| | | **F** | | | | | |
| DS26022409 (**7**) | 1-[(1-ethyl-3,5-dimethyl-1*H*-pyrazol-4-yl)methyl]-*N*-(4-isopropylphenyl)-**3-piperidinamine** | 354.5 | 4.73 | 4 | 1 | 3 | 33.1 |
| | | **G** | | | | | |
| DS26418355 (**8**) | (2,4-dimethoxyphenyl)(1-{[4-(4-fluorophenyl)-**1,3-thiazol-2-yl**]**methyl}-3-piperidinyl**) methanone | 440.5 | 4.37 | 7 | 6 | 0 | 79.9 |

| | | | | | | | |
|---|---|---|---|---|---|---|---|
| | | **C** | | | | | |
| DS27118552 (**9**) | 3-isopropyl-1-methyl-*N*-[(2-methyl-2,3-dihydro-1-benzofuran-5-yl)methyl]-1*H*-**pyrazole-5-carboxamide** | 313.4 | 3.19 | 4 | 1 | 3 | 56.2 |
| | | **Inhibitors** | | | | | |
| | | **C** | | | | | |
| DS72499364 (**10**) | 1-(4-methoxybenzyl)-*N*-methyl-*N*-(4-quinolinylmethyl)-1*H*-**1,2,3-triazole-4-carboxamide** | 387.4 | 3.16 | 6 | 0 | 5 | 73.1 |
| | | **D** | | | | | |
| DS15648900 (**11**) | methyl {[4-[1-(3-fluorobenzoyl)-**3-piperidinyl**]-5-(4-fluorophenyl)-2-pyrimidinyl]thio}acetate | 483.5 | 4.64 | 6 | 0 | 5 | 97.7 |
| DS27464324 (**12**) | [1-[(6-chloro-1,3-benzodioxol-5-yl)methyl]-3-(2-fluorobenzyl)-**3-piperidinyl**]methanol | 391.9 | 3.96 | 5 | 1 | 4 | 41.9 |
| DS38101551 (**13**) | [1-[(6-chloro-1,3-benzodioxol-5-yl)methyl]-3-(2-phenoxyethyl)-**3-piperidinyl**]methanol | 403.9 | 4.10 | 7 | 1 | 5 | 51.2 |
| | | **H** | | | | | |
| DS16594488 (**14**) | {1-[(4-ethylphenyl)**sulfonyl]-3-piperidinyl**}(3-methyl-2-thienyl)methanone | 377.5 | 4.92 | 3 | 0 | 4 | 91.1 |
| | | **I** | | | | | |
| DS24005623 (**15**) | *N*-(2,4-difluorobenzyl)-*N'*-[5-(1,1-dimethylpropyl)-1,3,4-**thiadiazol-2-yl]urea** | 340.4 | 4.29 | 5 | 2 | 3 | 95.2 |
| | | **J** | | | | | |
| DS48038642 (**16**) | (3S*,4S*)-1-(5-ethylpyrimidin-2-yl)-4-(2-naphthyl)**piperidin-3-ol** | 333.4 | 3.61 | 3 | 1 | 3 | 49.3 |
| | | **K** | | | | | |
| DS53041682 (**17**) | *N*-benzyl-7-(trifluoromethyl)-3,4-dihydroisoquinoline-2(1*H*)-**sulfonamide** | 370.4 | 4.50 | 2 | 1 | 2 | 57.8 |
| | | **L** | | | | | |
| DS54093530 (**18**) | *N*-{1-[2-(1-azocanyl)-**4-methyl-5-pyrimidinyl]ethyl**}-1-methyl-1*H*-pyrazole-**5-carboxamide** | 356.5 | 3.24 | 4 | 1 | 5 | 75.9 |

*DIVERSet Compound ID and Code
MW: Molecular Weight (g/mol)
clogP: calculated partition coefficient for lipophilicity: the affinity of the compound to a lipophilic environment
Rot. Bonds: non-H Rotatable bonds
Hdon: Hydrogen donor
HAcc: Hydrogen Acceptor
tPSA: total polar surface area
